# OligoArchive-DSM: Columnar Design for Error-Tolerant Database Archival using Synthetic DNA

**DOI:** 10.1101/2022.10.06.511077

**Authors:** Eugenio Marinelli, Yiqing Yan, Virginie Magnone, Marie-Charlotte Dumargne, Pascal Barbry, Thomas Heinis, Raja Appuswamy

## Abstract

The surge in demand for cost-effective, durable long-term archival media, coupled with density limitations of contemporary magnetic media, has resulted in synthetic DNA emerging as a promising new alternative. Today, the limiting factor for DNA-based data archival is the cost of writing (synthesis) and reading (sequencing) DNA. Newer techniques that reduce the cost often do so at the expense of reliability, as they introduce complex, technology-specific error patterns. In order to deal with such errors, it is important to design efficient pipelines that can carefully use redundancy to mask errors without amplifying overall cost. In this paper, we present OligoArchive-DSM (OA-DSM), an end-to-end DNA archival pipeline that can provide error-tolerant data storage at low read/write costs. Central to OA-DSM is a database-inspired columnar encoding technique that makes it possible to improve efficiency by enabling integrated decoding and consensus calling during data restoration.

## 1 INTRODUCTION

The global datasphere, or the sum total of all digital data generated, is expected to reach 125 Zettabytes by 2025, and over 50% of such data will be enterprise data stored in various databases, data lakes, and warehouses [9]. Today, over 80% of data generated is “cold”, or infrequently accessed, and corresponds to data that needs to be archived in order to meet safety, legal and regulatory compliance requirements [19]. Archival data is the fastest growing data segment with over 60% cumulative annual growth rate [26]. As enterprises continue migrate to the cloud, cloud vendors are in need of archival storage technologies that can provide high-density, low-cost storage of such data for decades without degradation. As all current storage media suffer from density scaling and durability limitations, researchers have started investigating radically new medium optimized for long-term archival. One such medium that has received a lot of attention recently is synthetic DNA.

DNA as a storage medium is seven orders of magnitude denser than tape [8] and can store up to 1 Exabyte of data in a cubic millimeter [7]. It is extremely durable and can last several millenia when stored under proper conditions. DNA is read by a process called sequencing, and the sequencing technology used to DNA is decoupled from DNA, the storage medium, itself. Thus, DNA will not suffer from obsolescence issues as we will always be able to read back data stored in DNA. Finally, using common, well-established biochemical techniques, it is very easy to replicate DNA rapidly. Thus, data stored in DNA can be easily copied. Given these benefits, several researchers have demonstrated the feasibility of using DNA as a long-term archival storage medium [1, 3, 6, 7, 10, 13, 15, 21, 25].

The primary obstacle to DNA storage adoption today is the prohibitive cost of reading and writing. The biochemical processes used for writing (*synthesis*) and reading (*sequencing*) DNA today were originally designed for biological applications that require very high precision and low scale. Using DNA as a storage medium requires a different trade off, as one can tolerate more errors in synthesis in sequencing for improved cost efficiency and scaling. In order to provide reliable data storage on DNA despite such errors, state-of-the-art (SOTA) approaches rely on using a significant amount of redundancy in both writing (in the form of parity bits generated by error control coding) and reading pipelines (in the form of very high sequencing coverage). The added redundancy, however, has the undesirable side effect of amplifying the read/write cost. Thus, efficient handling of errors is crucial to reducing overall cost.

In this work, we present *OligoArchive-DSM(OA-DSM)*, an end-to-end pipeline for DNA storage that provides substantially lower read/write costs than SOTA approaches. The core contributions of this work, and the two key aspects of OA-DSM that distinguish it from SOTA approaches are: (i) a novel, database-inspired, columnar encoding method for DNA storage, and (ii) an integrated consensus and decoding technique that exploits the columnar organization. In the rest of this paper, we provide an overview of challenges in DNA storage (Section 2), present the aforementioned aspects of OA-DSM design in detail (Section 3), and demonstrate their ability to achieve better accuracy and higher error-tolerance than SOTA methods using both simulation studies and a real wetlab validation experiment where we succesfully encoded and decoded a 1.2MB compressed TPC-H database archive.

## 2 BACKGROUND

Figure 1 provides an overview of SOTA DNA storage pipelines. Digital data is stored on DNA by first encoding bits into quaternary sequence of *nucleotides* (Adenine, Guanine, Cytosine, Thymine) – building blocks of the DNA macromolecule. These sequences are then used to create DNA molecules, or oligonucleotides (oligos), via a chemical process called *synthesis*. Data stored in DNA is read back via *sequencing*. Both synthesis and sequencing are approximate in nature and prone to errors. Thus, what is retrieved back from DNA are noisy copies of the original sequence referred to as *reads*. Thus, SOTA pipelines use consensus calling to infer original sequences from these reads. The inferred sequences are then decoded back into bits.

**Figure 1:**
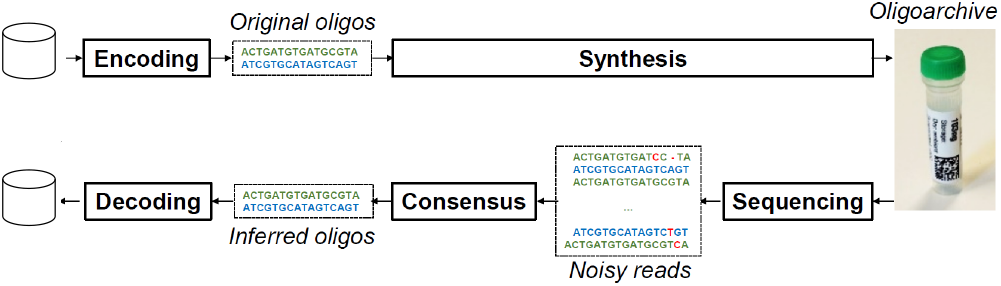
Read/write pipelines of SOTA DNA storage solutions.

There are several challenges in using DNA as a digital storage medium. First, not all DNA molecules can be synthesized or sequenced. There are several biological constraints (G-C constraint, homopolymer repeats, secondary structure formation, etc.) that must be respected during encoding to ensure downstream compatibility. Second, current synthesis processes cannot synthesize oligos longer than a few hundred nucleotides. Thus, as a single oligo can not store more than a few hundred bits at best, it is necessary to fragment the data and encode it across several oligos. Third, as DNA molecule itself has no addressing, it is necessary to add addressing information explicitly in the oligo during encoding in order to be able to reorder the oligos later during decoding. Fourth, as mentioned earlier, synthesis and sequencing are error prone. There can be insertion errors, where extra nucleotides are added to the original oligo resulting in a read being longer than the oligo, deletion errors where nucleotides are deleted resulting in shorter reads, and substitution errors. The error rates can vary depending on the technology used. Fifth, DNA storage also suffers from a coverage bias, where some oligos can be covered by multiple reads, and others can be completely missing (drop out). This happens due to physical redundancy during synthesis, where multiple DNA molecules are created for a single oligo, and uneven amplification of different oligos during sequencing library preparation (like Polymerase Chain Reaction that is used to amplify DNA before sequencing).

In order to ensure reliable data storage despite these errors, SOTA encoding methods rely on two distinct functionalities: (i) error control coding and (ii) consensus calling. During the write pipeline, input data bits are grouped into blocks (Fig 2(a)), and each block is encoded using error-correction codes, like Reed Solomon codes, LDPC, or fountain codes, to generate parity bits (Fig 2(b)). The original data and parity bits are then fragmented to divide them across oligos and indexed (Fig 2(b)). Finally, each indexed fragment is converted into an oligo (Fig 2(c)). During the read pipeline, the noisy reads that are produced by sequencing are fed to a consensus caller whose goal is to group similar reads and infer the original sequences. It is important to note here that these consensus sequences will not be error-free, accurate reproductions of original oligos. Instead, as the error rate and coverage bias increases, they will have errors and drop out. Hence, it is the job of the error-control decoder to use the additional parity bits to recover original input data despite these errors.

**Figure 2:**
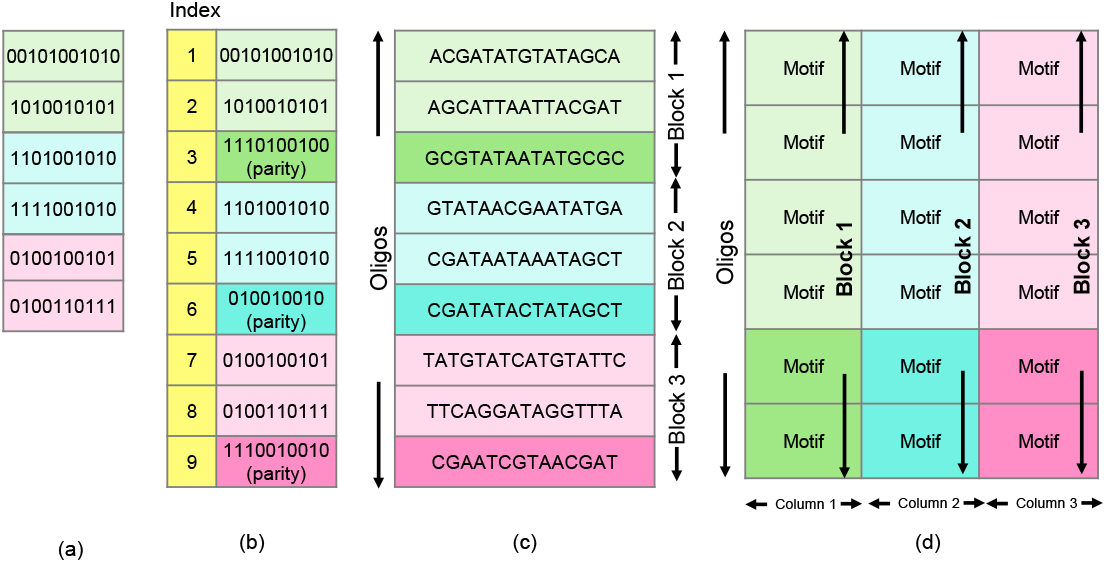
Comparison of SOTA versus OA-DSM columnar layout of oligos. The figures shows the raw input data being grouped into blocks (a), each block encoded to generate parity and indexed (b). (c) shows each block of input being mapped to multiple oligos with SOTA approaches. (d) shows each block being mapped to one column of motifs with OA-DSM.

To summarize, all SOTA pipelines share two characteristics: (i) an error-control-coded block of input data is encoded to generate a group of oligos that forms a unit of recovery, (ii) an isolated consensus step is performed before decoding to infer oligos from noisy reads. The decoding and consensus stages are independent steps in all SOTA pipelines and do not interact with each other.

## 3 DESIGN

Our approach to archiving data in DNA differs from SOTA based on the key observation that the separation of consensus and decoding is a direct side-effect of the data layout, that is the way oligos are encoded. Mapping a coded block of data to a group of oligos results in that group becoming a unit of recovery. Thus, before data can be decoded, the entire group of oligos must be reassembled by consensus, albeit with errors. The key idea in OA-DSM is to change the layout from the row-style SOTA layout (Figure 2(c)) to a database-inspired columnar layout (Figure 2(d)). A set of oligos is viewed like a relation, with each oligo being a row. OA-DSM encodes and decodes data one column at a time. The key benefit of this, as we show later in this section, is the fact that OA-DSM can integrate decoding and consensus into a single step, where the error-correction provided by decoding is used to improve consensus accuracy, and the improved accuracy in turn reduces the burden on decoding, thereby providing a synergistic effect. In the rest of this section, we will explain the OA-DSM design in more detail by presenting its read and write pipelines.

### 3.1 OA-DSM Write Pipeline

The top half of the Figure 3 shows the OA-DSM data writing pipeline. The input to the write pipeline is a stream of bits. Thus, any binary file can be stored using this pipeline. The first step in processing the input involves grouping it into blocks of size 256,000 bits. Each block of input is then randomized. While it is not relevant to this discussion, we use randomization similar to SOTA to improve the accuracy of read clustering in the data decoding stage as explained in Section 3.2. After randomization, error correction encoding is applied to protect the data against errors. We use Low-Density Parity Check (LDPC) codes [12] with a block size of 256,000 bits. Prior work has demonstrated that such a large-block-length LDPC code is resilient to both substitution/indel errors, that cause reads to be noisy copies of original oligos, and synthesis/sequencing-bias-induced dropout errors, where entire oligos can be missing in reads due to lack of coverage [6].

**Figure 3:**
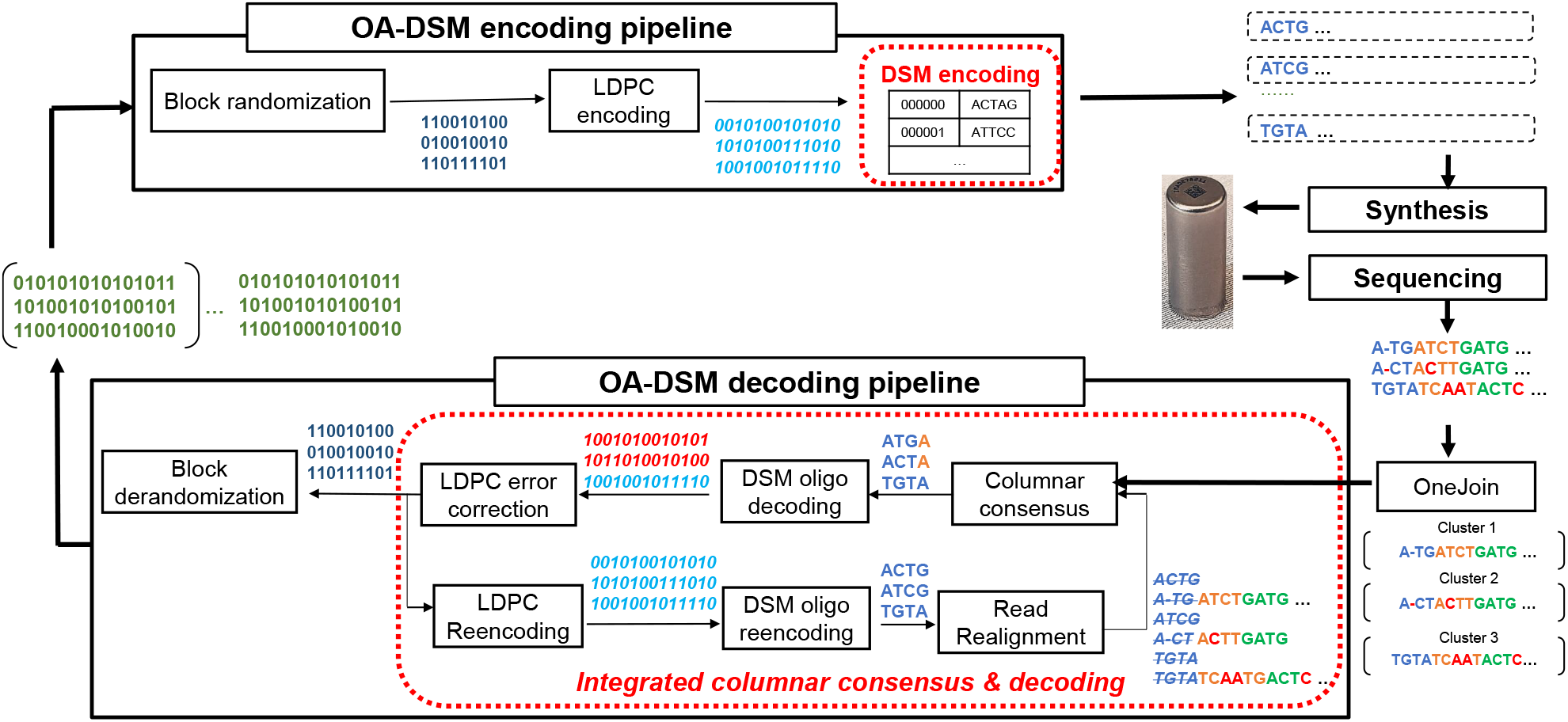
OA-DSM data writing (top) and reading (bottom) pipelines. The blocks in red are unique to OA-DSM (versus SOTA).

The LDPC encoded bit sequence is fed as input to the *DSM-oligo-encoder* which converts bits into oligos. While SOTA approaches design each oligo as a random collection of nucleotides, the DSM-oligo-encoder designs oligos using composable building blocks called *motifs*. Each motif is itself a short oligo that obeys all the biological constraints enforced by synthesis and sequencing. Multiple motifs are grouped together to form a single oligo. We use motifs rather than single nucleotides as building blocks because, as we will see later in Section 3.2, integration of decoding and consensus relies on alignment which cannot be done over single nucleotides.

In order to perform the conversion of bits into motifs, the DSM-oligo-encoder maintains an associative array with a 30-bit integer key and a 16 nucleotide-length (nt) motif value. This array is built by enumerating all possible motifs of length 16nt (AAA, AAT, AAC, AAG, AGA…) and eliminating motifs that fail to meet a given set of biological constraints. We configure our encoder to admit motifs that have up to two homopolymer repeats (AA,CC,GG, or TT), and GC content in the range 0.25 to 0.75. With these constraints, using 16nt motifs, out of 4^16^ possible motifs, we end up with 1,405,798,178 that are valid. By mapping each motif to an integer in the range 0 to 2^30^, we can encode 30-bits of data per motif. Thus, at the motif level, the encoding density is 1.875 bits/nt. While we can increase this density by increasing motif size, or relaxing biological constraints, we limited ourselves to this configuration due to two reasons: (i) memory limitation of our current hardware, as the current associative array itself occupies 100GB of memory, (ii) the motif design is orthogonal to the columnar encoding which is the focus of this work. In future work, we plan to increase this bit density by expanding to large motif sets.

The use of motifs as building blocks renders a distinct relational organization to oligos–just as a set of attributes form a tuple, and a set of tuples form a relation, a set of motifs (attributes) forms an oligo (row), and a set of oligos constitutes an OligoArchive. Thus, the second major difference of our approach to SOTA is the layout of motifs across oligos which is reminiscent of Decomposition Storage Model (DSM), or columnar data layout, adopted by modern analytical database engines. The motifs generated from an error-control coded data block are used to extend oligos by adding a new column as shown in Figure 2(d). This process is repeated until the oligos reach a configurable number of columns after which the process is reset to generate the next batch of oligos again from the first column. The generated oligos can then be synthesized to produce DNA molecules that archive data.

While the figure shows all columns as being of the same size, a small subtlety in the practical implementation is the distinction between the first column and the rest. As we need to index the oligos to enable reordering during decoding, the first column of motifs is generated by using a 15-bit address and a 15-bit data to generate a 30-bit integer. Thus, the first LDPC encoded block is decomposed into 15-bit integers. However, from the second column, there is no need to add addressing information. Thus, rest of the LDPC blocks are decomposed into 30-bit integers. As a result, all columns except the first encode 2 LDPC blocks, while the first column encodes only 1 LDPC block. Note that with 15-bit addresses, we can address up to 32,768 oligos. In ongoing work, we are using OA-DSM to develop a block-addressed, randomly-accessible, DNA file system. Similar to traditional file systems, OA-DSM allows us to view a column like a disk block, and a collection of columns like an extent. The 15-bit address here provides intra-extent addressing. Extents themselves will be addressed separately using a separate mechanism (nested primers). We explicitly mention this here to clarify that OA-DSM can scale to much larger oligo pools. But for the rest of this paper, we focus on columnar design and consensus.

### 3.2 OA-DSM Read Pipeline

As mentioned before, data stored in DNA is read back by sequencing the DNA to produce reads, which are noisy copies of the original oligos that can contain insertion, deletion, or substitution errors. As each oligo can be covered by multiple reads, the first in decoding is clustering to group related reads together. In prior work, we developed an efficient clustering technique based on edit similarity joins [22, 23] that exploits the fact that due to randomization during encoding, reads corresponding to the same original oligo are “close” to each other despite errors and “far” from the reads related to other oligos. The output of this algorithm is a set of clusters, each corresponding to some unknown original oligo.

After the clustering stage, other SOTA methods apply consensus in each cluster followed by decoding in two separate phases as shown in Figure 1. In OA-DSM, we exploit the motif design and columnar layout of oligos to iteratively perform consensus and decoding in an integrated fashion as shown in Figure 3. Unlike other approaches, OA-DSM processes the reads one column at a time. Thus, the first step is columnar consensus which takes as input the set of reads and produces one column of motifs. The choice of consensus algorithm is orthogonal to OA-DSM design. We use an alignment-based bitwise majority algorithm we developed previously for consensus [23], as we found this to provide accuracy comparable to other state-of-the-art trace reconstruction solutions [2]. The motifs obtained from consensus are then fed to the *DSM-oligo-decoder* which is the inverse of the encoder, as it maps the motifs into their 30-bit values. Note here that despite consensus, the inferred motifs can still have errors. These wrong motifs will result in wrong 30-bit values. These errors are fixed by the *LDPC-decoder*, which takes as input the 30-bit values corresponding to one LDPC block and produces as output the error-corrected, randomized input bits. These input bits are then derandomized to produce the original input bits for that block.

As mentioned earlier, SOTA methods do not use the error-corrected input bits during decoding. OA-DSM, in contrast, uses these bits to improve accuracy as shown in the bottom part of the integrated columnar consensus in Figure 3. The error-corrected bits produced by the LDPC-decoder are reencoded again by passing them through the LDPC-encoder and DSM-oligo-encoder. This once again produces a column of motifs as it would have been done during input processing. The correct column of motifs is used to realign reads so that the next round of columnar decoding starts at the correct offset. The intuition behind this realignment is as follows. An insertion or deletion error in the consensus motifs will not only affect that motif, but also all downstream motifs also due to a variation in length. For instance, if we look at the example in Figure 3, we see a deletion error in read *A — TATCTG..* which should have been *ATCGATCTG* This results in the first motif being incorrectly interpreted as *ACTA* (instead of *ACTG*, and second motif as *TCTG* (instead of *ATCT*). Thus, an error early in consensus keeps propagating. Without a knowledge of the correct motif, there is no way to fix this error. But in OA-DSM, by reencoding the error-corrected bits, we get the correct motifs. By aligning these motifs against the reads, we can ensure that consensus errors do not propagate. Note here that such realignment is only possible because we use motifs, as two sequences can be aligned accurately only if they are long enough to identify similar subsequences. Thus, columnar layout without motifs, or with just nucleotides, would not make realignment possible. Similarly, integrating consensus and decoding is possible only because of the columnar layout, as the SOTA layout that spreads a LDPC block across several oligos cannot provide incremental reconstruction.

## 4 EVALUATION

In this section, we will present the results from our experimental evaluation of the OA-DSM pipeline. The evaluation is structured as follows. First, we show the advantage of using a columnar design by comparing OA-DSM with a row-based pipeline (Sec. 4.1). Then, we compare OA-DSM with various SOTA approaches with respect to read cost and write cost to show that our design can lead to substantial cost reduction (Sec. 4.2). Finally, we present preliminary results from ongoing wetlab experiments to validate the end-to-end OA-DSM pipeline (Sec. 4.3).

We conduct all the experiments on a local server equipped with a 12-core CPU Intel(R) Core(TM) i9-10920X clocked at 3.50GHz, 128GB of RAM. The core components of the OA-DSM pipeline shown in Figure 3 has been implemented in C++17. We use TPC-H dbgen utility to generate compressed, synthetic data that we treat as the archival file that must be stored on DNA. We parameterize dbgen to control the generated database size according to experimental requirements as mentioned later.

### 4.1 Benefits of Columnar Design

In order to ensure that the benefits of OA-DSM are due to the columnar design and not other parameters, we have developed a row-based version of the pipeline shown in Figure 3, where we fixed all other parameters (clustering and consensus algorithms, LDPC block size, motif set, etcetera), and only changed two aspects to make it similar to SOTA: (i) replace DSM encoder with row-based encoder that maps one LDPC block to multiple oligos, (ii) perform consensus to infer entire oligos first, and then decode separately.

In order to compare the columnar and row-based pipelines, we perform an end-to-end DNA storage simulation study using both pipelines. First, we use both pipelines to generate the oligos for a 3MB TPC-H archive file (3MB size was chosen based on calculations that ensure that both pipelines produce the same number of oligos). We configure LDPC encoder to generate two datasets, with 10% and 30% redundancy. Then, we encode the two datasets using both pipelines, while fixing the oligo length to 50 motifs per oligo (800nt), generating four OligoArchives, two containing 18773 oligos (row/column at 10% redundancy), and the other two containing 22187 (row/column at 30% redundancy) oligos.

We compare the row and columnar pipelines by evaluating the minimum coverage required at 10% and 30% redundancy levels to achieve 100% error-free reconstruction of the input data at various the error rates (1% to 12%). We conduct the experiment similar to SOTA [6, 21] as follows. For each error rate, and for each of the four oligo sets, we generate read datasets at various coverage levels (1 to 25). In order to generate reads, we first duplicate each oligo a certain number of times according to the configured coverage level. Then we inject random errors at random positions in each read. We inject insertion, deletion and substitution with an equal probability, and the number of errors injected per read follows a normal distribution with mean set to the configured error rate. We then decode the read datasets using both pipelines and identify the minimum coverage level required to fully recover the original data.

Figure 4 shows the minimum coverage for data encoded with 10% redundancy. Clearly, columnar-wise encoding outperforms the row-wise one, as it reduces the coverage required up to 40% for high error rates. This reduction in minimum coverage can be intuitively explained as follows. Row-based encoding maps an LDPC block into multiple oligos. This implies that a single erroneous oligo can lead to a data loss of up to 1500 bits (50 motifs per oligo x 30 bits per motif). As explained in Section 3, all that is required for an oligo loss is a single insertion/deletion error in the first motif after consensus. On the other hand, an oligo loss in OA-DSM only causes a loss of 30 bits in each of the LDPC blocks, thanks to the columnar encoding. Further, the integrated consensus and decoding can fix consensus errors in early rounds so that they do not affect future rounds. Due to these reasons, the LDPC decoder works much more effectively when paired with columnar layout rather than row-based encoding. The results are similar for data encoded with 30% redundancy as well, as shown in Figure 5. Notice that in the 30% case, both row and OA-DSM pipelines have a minimum coverage lower than the 10% redundancy case. This is expected, as a higher redundancy implies a higher tolerance to errors.

**Figure 4:**
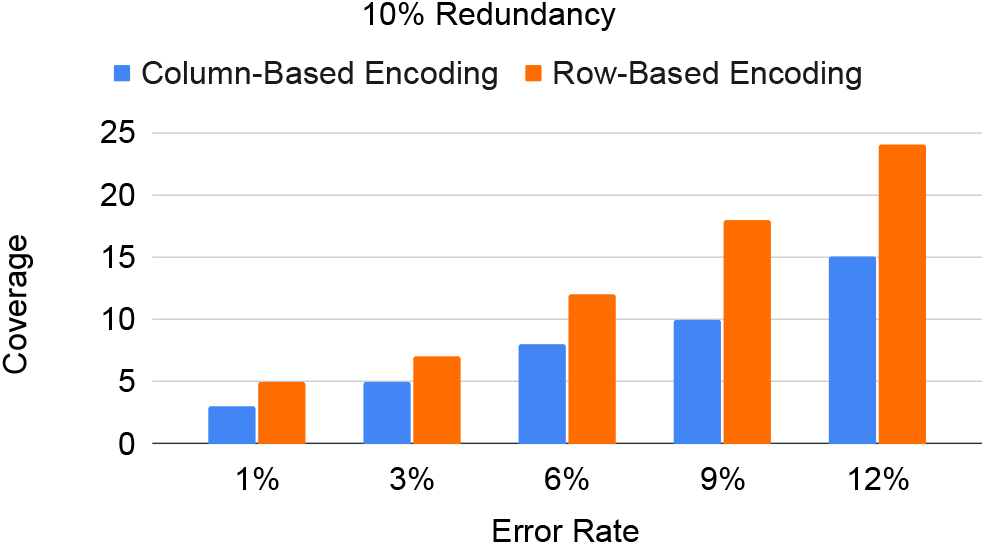
Min. cov. at 10% redundancy.

**Figure 5:**
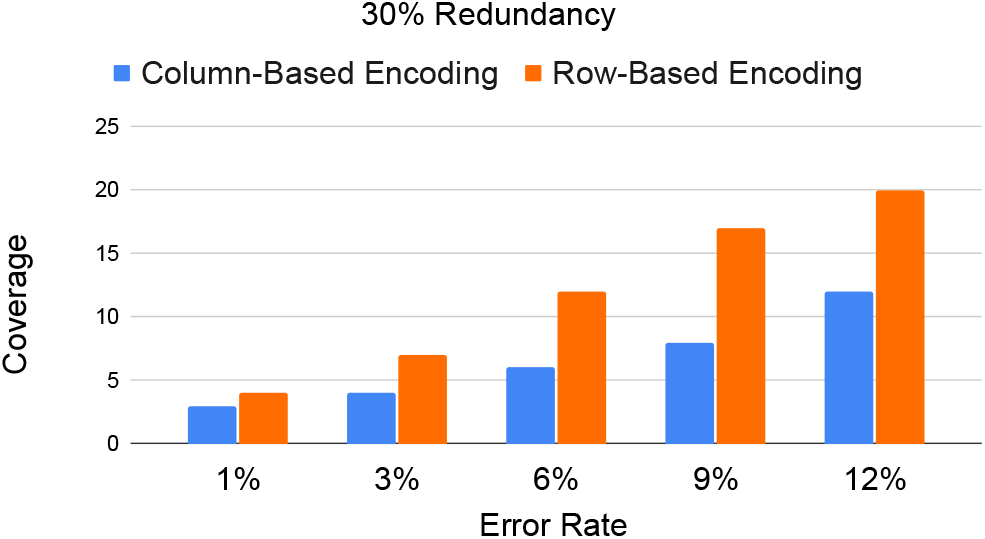
Min. cov. at 30% redundancy.

### 4.2 SOTA comparison

Having demonstrated the advantage of using a columnar storage architecture, we will now present a comparison of OA-DSM with SOTA approaches in terms of reading and writing cost [6, 21, 25]. Writing cost is defined as 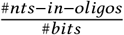, where the numerator is the product of the number of oligos and the oligo length, and the denominator is the input data size. Thus, higher the redundancy and encoding overhead, higher the write cost. The reading cost is defined by 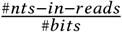. The numerator is the sum total of all read lengths, and denominator is the input size. Thus, higher the coverage required, higher the read cost.

Figure 6 shows the read and write cost for OA-DSM and other SOTA algorithms. For OA-DSM, we compute these costs based on results shown in Figures 4 and 5. For SOTA approaches, we reproduce the costs from their publications. There are several observations to be made. First, let us compare the OA-DSM with row-based SOTA approach that also uses LDPC (by S. Chandak et al. [6]). Both these cases use the same LDPC encoder configured with 30% redundancy. The cost reported here is for 1% error rate in both cases. Clearly, the OA-DSM approach has both a lower write and read cost. The difference in write cost can be explained due to the fact that in the row-based LDPC approach, the authors also added additional redundancy in each oligo in the form of markers which they used in their decoder. OA-DSM is able to achieve 100% data reconstruction using the same LDPC encoder at a much lower coverage level without such markers as demonstrated by the lower read cost.

**Figure 6:**
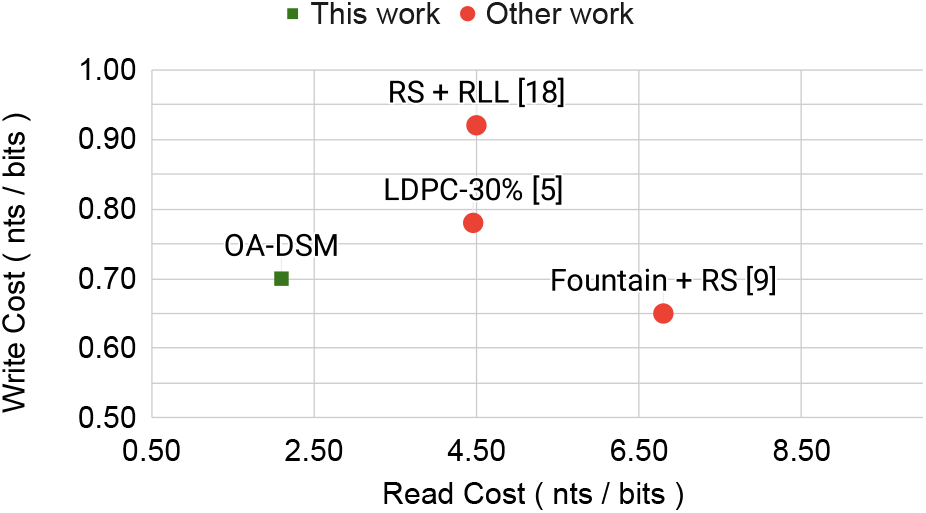
OA-DSM vs. SOTA rd/wt costs.

Comparing OA-DSM with the other two efficient encoders (large-block Reed-Solomon coding by Organick et al. [25] and fountain codes by Erlich et al. [10]), we see that OA-DSM provides substantially better read cost, but slightly worse write cost than fountain coding approach. As we mentioned earlier, we can further improve the write cost for OA-DSM using several approaches. First, the OA-DSM results in Figure 6 were obtained with a 30% redundancy based on its ability to handle even 12% error rate. For lower error rates (less than 1%), as was the case with the Fountain coding work, even 10% redundancy would be able to fully restore data at extremely low coverage (3× as shown in Figure 4). Second, as mentioned in Section 3, scaling the motif set by using longer motifs (17nt and 33 bits) could allow us to increase bit-level density further from 1.87 bits/nt to over 1.9 bits/nt. These two changes would lead to further reduction in write cost without any adverse effect on the read cost. As this work was predominantly about reducing the read cost, we leave open these optimizations to future work.

Finally, Lin et al. [21] recently presented the Gini architecture which interleaves nucleotides across oligos in order to minimize the impact of consensus errors. We also tried to compare OA-DSM with Gini, but we could not derive the read/write cost for Gini, which was also not reported, due to lack of statistics about reads. However, as our evaluation methodology is identical to Gini, we present a direct comparison of results in terms of minimum coverage required by both approaches.Figure 7 shows the minimum coverage required by OA-DSM, Gini, and a baseline without Gini reported by Lin et al. [21], to perfectly recover data at various error rates. At 18.4% redundancy based on Reed-Solomon coding, the reported baseline needed 30× to recover data 12% error rate. Gini, in contrast, provided a 33% improvement as it needed a minimum coverage of 20× at 12% error rate to guarantee full recovery. OA-DSM configured at 30% redundancy with LDPC encoding provides a 40% improvement over Gini, it requires only 12× coverage. Comparing Figure 7 with Figure 4, we see that OA-DSM provides 25% less coverage (15×) even at 10% redundancy compared to Gini. Thus, OA-DSM has a much lower read cost, thanks to the integrated consensus and decoding enabled by columnar organization.

**Figure 7:**
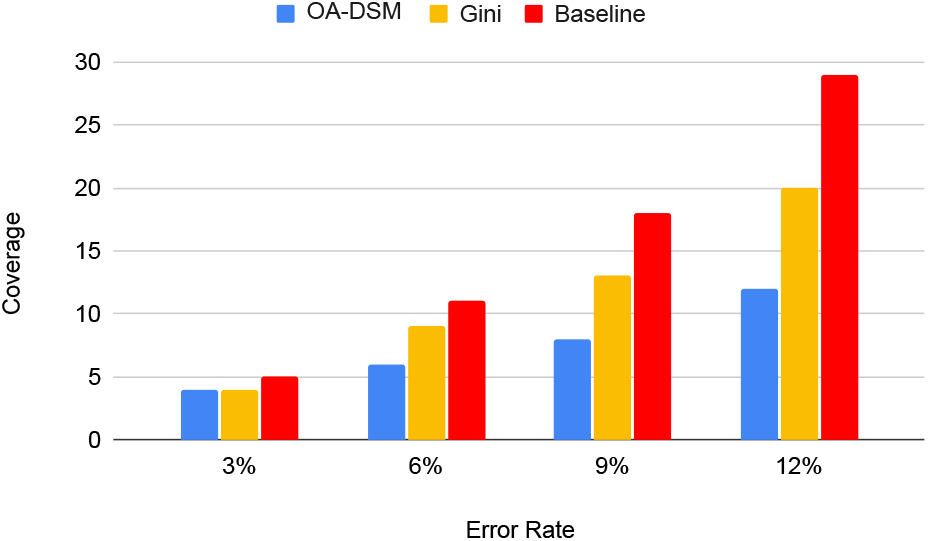
Comparison to Gini.

### 4.3 Wetlab validation

Finally, we present the details of our ongoing wetlab validation experiment, in which we are storing a TPC-H database compressed in a single archive file of 1.2MB. We limited the size to 1.2MB to limit the cost of actual synthesis. The cardinality of various relations in the archive is reported in Table 1. Using OA-DSM configured with 30% LDPC redundancy, we encoded the archive file to generate 44376 oligos, with each oligo of length 160nts (length chosen to optimize synthesis cost). The oligos were synthesized by Twist Biosciences for a cost of approximately €10,000. We sequence the synthesized oligos using Oxford Nanopore PromethION platform generating approximately 43 millions of noisy reads.

**Table 1:** TPCH database summary for WetLab experiment.

| Table Name | Number of Rows | Size [Byte] |
| --- | --- | --- |
| customer | 900 | 142782 |
| lineitem | 31220 | 3717358 |
| nation | 25 | 2199 |
| orders | 9000 | 982200 |
| partsupp | 4800 | 690415 |
| part | 1200 | 140596 |
| region | 5 | 384 |
| supplier | 60 | 8242 |

To perform error characterization, we aligned the 43M reads to the original oligos using BWA-MEM seqence aligner[20]. 99.99% reads were aligned to a reference oligo, indicating a very high quality of the generated read set. Figure 8 shows the coverage histogram (number of oligos that have a given coverage). Each reference oligo is covered by at least one read, with a median coverage of 951×, minimum coverage of 8×, and a maximum coverage of 2500×. We deliberately sequenced the oligos at such high coverage to test recovery at various coverage levels as we present later.

**Figure 8:**
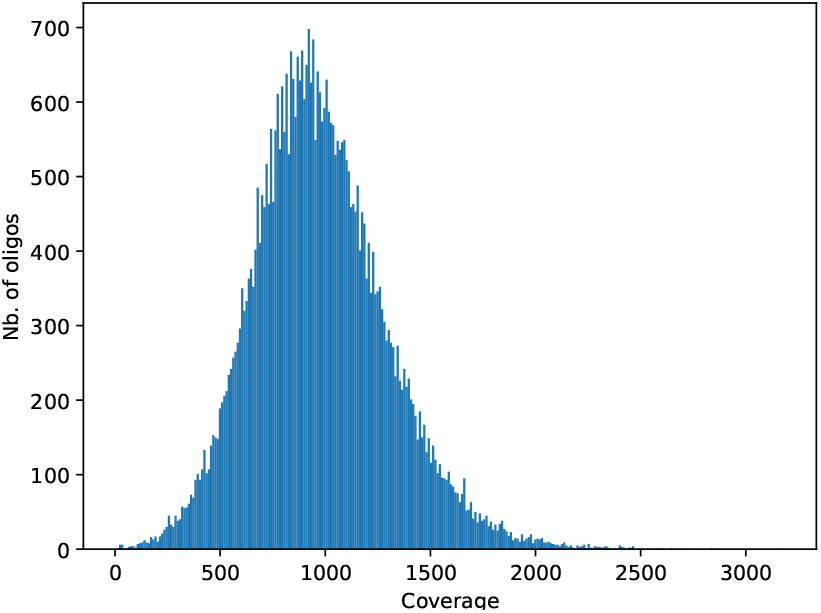
The histogram of oligos’ coverage.

Figure 9 shows the substitution, insertion and deletion rate per position (as computed by BBmap [5]). Note that while the data-carrying oligo had a length of 160, our reads are longer as they include the primers that were appended at both ends of the oligo for sequencing. As these primers get trimmed out during read preprocessing, the error rate of relevance to us is the middle portion of the read which corresponds to the encoded, data-carrying portion of the oligo. We see that in this portion, the substition rate is dominant, which is 3× higher than insertion and deletion rates. Figure 11 compares our error rates with those reported in prior work on DNA storage [11, 14, 16, 17, 24]. While the actual rates vary due to differences in synthesis and sequencing steps, we see that the overall trends are similar. Using the aligned reads, we also report the indel distribution in Figure 10 which shows a histogram of edit distances between the reads and references. As can be seen, 96.97% reads have edit distance less than 10, indicating that the error rate is less than 6%.

**Figure 9:**
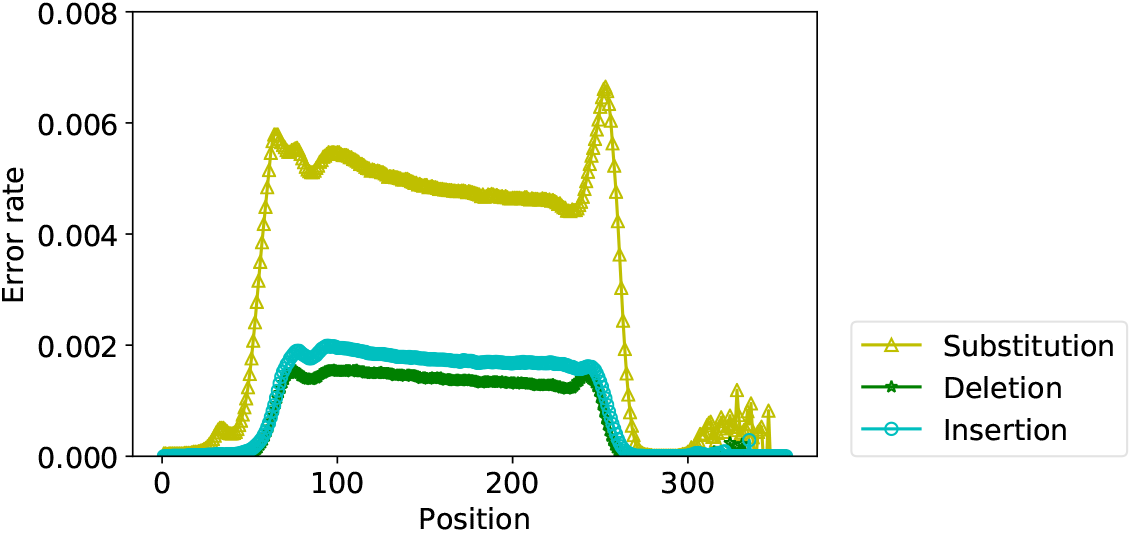
The substitution, insertion and deletion rate per position.

**Figure 10:**
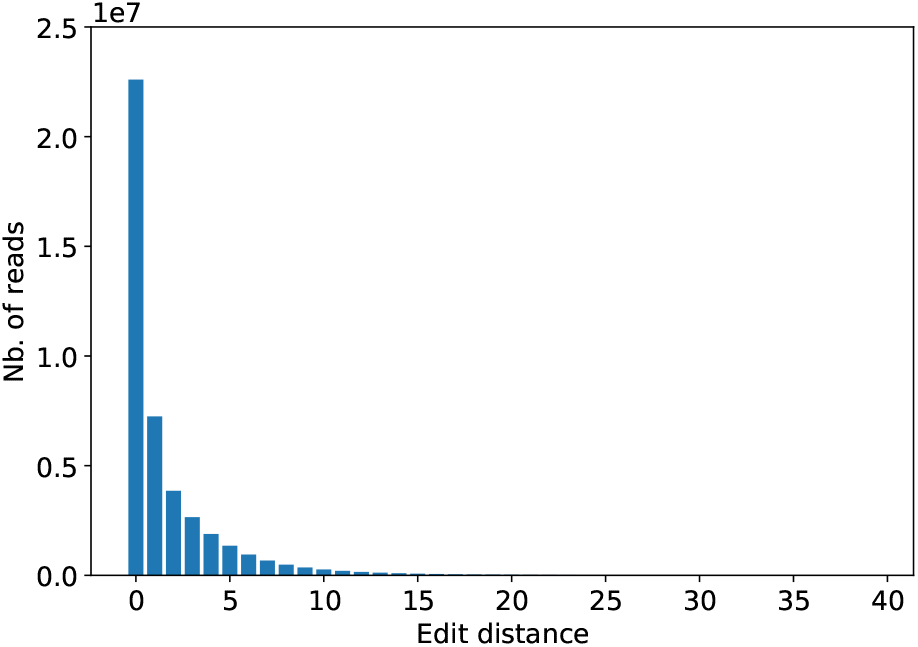
Distribution of edit distance.

**Figure 11:**
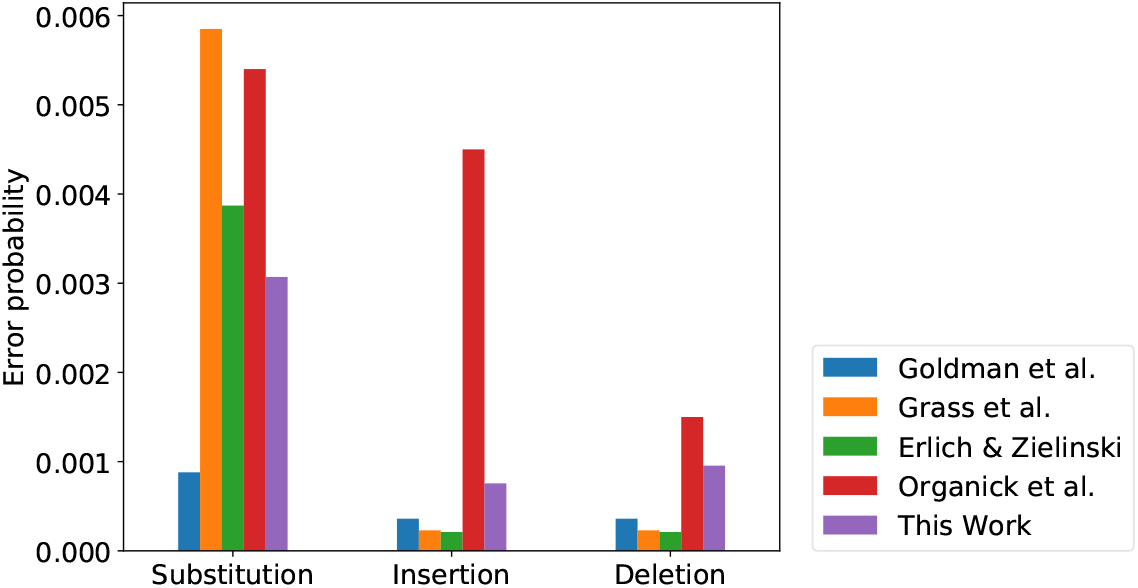
Comparison of errors with previous work.

In order to test end-to-end decoding, we first used the full 43M read dataset as input to the decoding pipline. Unsurprisingly, we were able to achieve full data reconstruction, given the ability of OA-DSM to handle much lower coverage levels and higher error rates. In order to stress test our decoding pipeline and identify the minimum coverage that allows fully reconstruction of data, we repeated the decoding experiment on smaller readsets which were derived by randomly sampling a fraction of reads from the 43M read dataset. In doing so, we found that OA-DSM was able to perform full recovery using just 200K reads, which corresponds to a coverage of 4×. At this coverage, nearly 3500 out of 44376 reference oligos were completely missing. However, the LDPC code and columnar decoding were able to successfully recover data. As further reduction in coverage led to data loss, we validate 4× as the minimum coverage OA-DSM can handle with our wetlab experiment. Computing the costs for minimum coverage, we get a read cost of 3.37 nts/bit, and a write cost of 0.72 nts/bit, similar to the results reported in Figure 6.

## 5 RELATED WORK

Biologists have long demonstrated the ability to store data in DNA [27]. However, using DNA as a “large-scale” data storage media is an entirely new area of research that has become possible only recently, thanks to advances in sequencing and synthesis technologies. Almost all prior work on DNA data storage has focused on developing encoding techniques for mapping binary files to oligos. We only present an overview of a few key related publications in this section. A comprehensive survey of approaches can be found here [18].

Pioneering work by Church et al. [7] was the first to prove the viability of DNA as a large-scale digital storage medium. Using a simple coding scheme that maps a 0 bit in the input stream to an A or C and a 1 bit to G or T, they stored a HTML file of 0.65MB. However, due to errors in sequencing and synthesis procedures, data retrieval was not fully automated and required manual intervention. Following this, Goldman et al. proposed a variable-length coding scheme that improves storage density by compressing and transforming data into an intermediate base-3 representation [13]. Using this procedure, they stored 739KB of digital data and showed that encoding/decoding of data can be done automated despite errors.

Follow up work investigated error-correction techniques that can provide full recovery while substantially improving storage density. Bornholt et al. optimized Goldman et al.’s strategy by implementing simple XOR-based redundancy [4]. Grass et al. [15] proposed an error-correcting scheme that used concatenation codes with Reed Solomon codes as both inner and outer codes to store 83KB of data. Blawat et al. [3] study the error properties of DNA storage based on the results from the first study by Church et al. Using a forward error-correction code customized to the DNA erasure channel, they stored 22MB of data. Organick et al. [25] propose a new concatenation code with Reed-Solomon as the outer code and the differential code from Goldman et al. as the inner code. Using this new code, they stored and recovered 200MB of data. We compared OA-DSM with several of these methods in Section 4.

## 6 CONCLUSION

All SOTA approaches for DNA data archival use a “row-based” approach for mapping input bits onto oligos. In this paper, we showed how this approach results in a strict separation of consensus calling and decoding, and how this separation, in turn, results in lost opportunity for improving read/write cost. We presented OA-DSM, an end-to-end pipeline for DNA data archival that uses a novel, database-inspired, columnar data organization. We showed how such an approach enables the integration of consensus and decoding stages so that errors fixed by decoding can improve consensus and vice versa. Using a full system evaluation, we highlighted the benefit of our design and showed that OA-DSM can substantially reduce read-write costs compared to SOTA approaches.

## Notes

### Competing Interest Statement

The authors have declared no competing interest.

